# Dissociable mechanisms of reward learning co-mature during human adolescence as predicted by macaque lesion models

**DOI:** 10.1101/2021.01.14.426647

**Authors:** Marco K. Wittmann, Maximilian Scheuplein, Sophie G. Gibbons, MaryAnn P. Noonan

**Author notes:** **Address for correspondence:** Department of Experimental Psychology, University of Oxford, Radcliffe Observatory, Anna Watts Building, Woodstock Rd, Oxford OX2 6GG. authors contributed equally to this work (Marco K. Wittmann, Maximilian Scheuplein, MaryAnn P. Noonan).

## Abstract

Reward-guided learning and decision-making is a fundamental adaptive ability and depends on a number of component processes. We investigate how such component processes mature during human adolescence. Our approach was guided by analyses of the effects of lateral orbitofrontal lesions in macaque monkeys, as this part of the brain shows clear developmental maturation in humans during adolescence. Using matched tasks and analyses in humans (n=388, 11-35yrs), we observe developmental changes in two key learning mechanisms as predicted from the monkey data. First, choice-reward credit assignment – the ability to link a specific outcome to a specific choice – is reduced in adolescents. Second, the effects of the global reward state – how good the environment is overall recently − exerts a distinctive pattern of influence on learning in humans compared to other primates and across adolescence this pattern becomes more pronounced. Both mechanisms were correlated across participants suggesting that associative learning of correct reward assignments and GRS based learning constitute two complementary mechanisms of reward-learning that co-mature during adolescence.

## Introduction

To achieve a desirable goal, humans and other animals need to learn about the consequences of their choices and chose rationally between them. Specifically, actions that lead to positive outcomes should be repeated in the future, while actions that do not result in rewards should be omitted (Rescorla & Wagner, 1972). This requires the formation of contingencies between choice options and outcomes, i.e. the assignment of credit for outcomes to choices (Thorndike, 1911; 1933a). However, specific deviations from appropriate choice-reward credit assignments and rational choice (Noonan *et al*., 2010; Noonan *et al*., 2017) exist. For example, when reward reinforces not just a single choice but multiple (Noonan *et al*., 2010; Walton *et al*., 2010; Jocham *et al*., 2016; Noonan *et al*., 2017; Rudebeck *et al*., 2017; Wittmann *et al*., 2020). We refer to this as non-contingent credit assignment. Despite the ubiquity of contingent and non-contingent credit in learning and decision-making processes, very little is known about how these subcomponents of reward learning mature (Hartley & Somerville, 2015). Understanding if and how these processes change during adolescence is particularly critical as this dynamic maturational period in life offers a window of opportunity during which interventions, such as those to reduce risk-taking, may have critical impacts on the developmental trajectory (Dahl *et al*., 2018).

During adolescence young people gain independence and develop mature social goals (Fuhrmann *et al*., 2015; van Duijvenvoorde *et al*., 2016), yet it is also a time characterised by irrational, risky and impulsive decisions, including drug-use (Blakemore & Robbins, 2012; Viner *et al*., 2012). Reward-related computations change during adolescence as corresponding neural substrates continue to develop (Galvan & McGlennen, 2013; Barkley-Levenson & Galvan, 2014; Insel *et al*., 2017). The orbitofrontal cortex is one of the last areas of the brain to mature, still showing signs of grey matter development into the early twenties, and it appears that the boundary region of lateral orbitofrontal, insular and ventrolateral prefrontal cortex may be particularly developmentally delayed (Gogtay *et al*., 2004). From this, we would predict developmental maturation during adolescence of reward-related computations that depend on this lateral prefrontal region.

Lesion studies have shown that damage to these lateral prefrontal regions lead to deficits in contingent credit assignment (Rudebeck *et al*., 2013; Noonan *et al*., 2017). While more recently, a separate part of this region has been shown to hold a representation of the global reward state (GRS) – how rewarding the environment is overall in the recent past (Wittmann *et al*., 2020). In a high GRS, macaque monkeys are more likely to stay with a rewarded option while they are more likely to abandon unrewarded choices if the GRS is low. This means that both contingent rewards that coincide with a choice, but also the GRS *per se*, increased the likelihood of repeating that choice. Because of these anatomical considerations, we hypothesized that both types of credit assignment undergo developmental changes during adolescence. By contrast, if the more medial orbitofrontal regions mature faster than the lateral regions, as visually suggested by current developmental maps (Gogtay *et al*., 2004), then we hypothesize that decision-making processes linked to these regions may not show similar developmental changes during adolescence. We test this hypothesis by examining a type of decision-making bias that occurs in multi-option environments and depends on medial prefrontal cortex: how the comparison of two option is influenced by an irrelevant third alternative (Ray, 1973; Noonan *et al*., 2010; Louie *et al*., 2011; Louie *et al*., 2013; Chau *et al*., 2014; Noonan *et al*., 2017; Chau *et al*., 2020).

Here, we draw on a multi-option probabilistic learning task first established in macaque monkeys that allows us to disentangle the above-mentioned subcomponents of reward learning and decision-making during human adolescence (Noonan *et al*., 2010; Walton *et al*., 2010; Chau *et al*., 2015; Noonan *et al*., 2017; Rudebeck *et al*., 2017). We tested a large online sample (overall *n*=422) of adolescents (age 11-17) and young adults (age 18-35) and delineate the developmental trajectory of these component processes. Importantly, our analyses of human developmental changes are guided by a reanalysis of macaque behavior during a matched probabilistic learning task (Noonan *et al*., 2010; Walton *et al*., 2010) using identical analysis approaches. We demonstrate that both contingent learning and learning based on the GRS is altered in macaques with lateral prefrontal lesions supporting the idea that these component mechanisms may continue to change in humans as lateral prefrontal cortex matures during adolescence (Gogtay *et al*., 2004; Raznahan *et al*., 2010). Indeed, we find both learning mechanisms change significantly during adolescence. Strikingly, we find a qualitative difference in the way both species rely on the GRS during learning suggesting that while the GRS may dilute optimal credit assignment in macaques, it may aide optimal switching behaviour in humans in line with some theoretical considerations on contrast effect in valuation (Crespi, 1942; Daw & Touretzky, 2002; McNamara *et al*., 2013). Consistent with this idea, the strength of the GRS effects correlated with the strength of contingent learning effects across human participants. By contrast, the biasing effects of irrelevant alternatives linked to medial prefrontal cortex remained unchanged across age groups. Together, this suggests that some subcomponents of reward learning in adolescence co-mature leading to overall less explorative, and more exploitative, patterns of learning and choice (Trudel et al., 2020; Wilson et al., 2014).

## Results

### Macaque lateral prefrontal lesions change the impact of contingent rewards and the GRS on choice

First, we analyzed data from seven macaques (4-10yrs) to approach the hypothesis that the mechanisms of reward learning that change during adolescence in humans can be understood in reference to neural investigations of the same learning processes in monkeys. Specifically, we considered lateral orbitofrontal cortex because this brain region supports credit assignments in animal models and continues to mature during adolescence (Gogtay *et al*., 2004). Therefore, we would expect subcomponents of reward learning that rely on this area to also exhibit measurable changes during this period in life in humans. We tested this approach empirically for the case of contingent credit assignment and learning from the GRS and contrast it with computations linked to more medial parts of prefrontal cortex. We analyzed macaque data from an experimental paradigm (Fig.1) nearly identical to our human experiment using the same regression model in both cases. This precise mapping of experimental paradigm and analysis technique across species allowed us to directly compare the mechanisms via which human adolescents learn with the way macaques learn from reward under normal circumstances and after lateral lesions.

**Figure 1.**
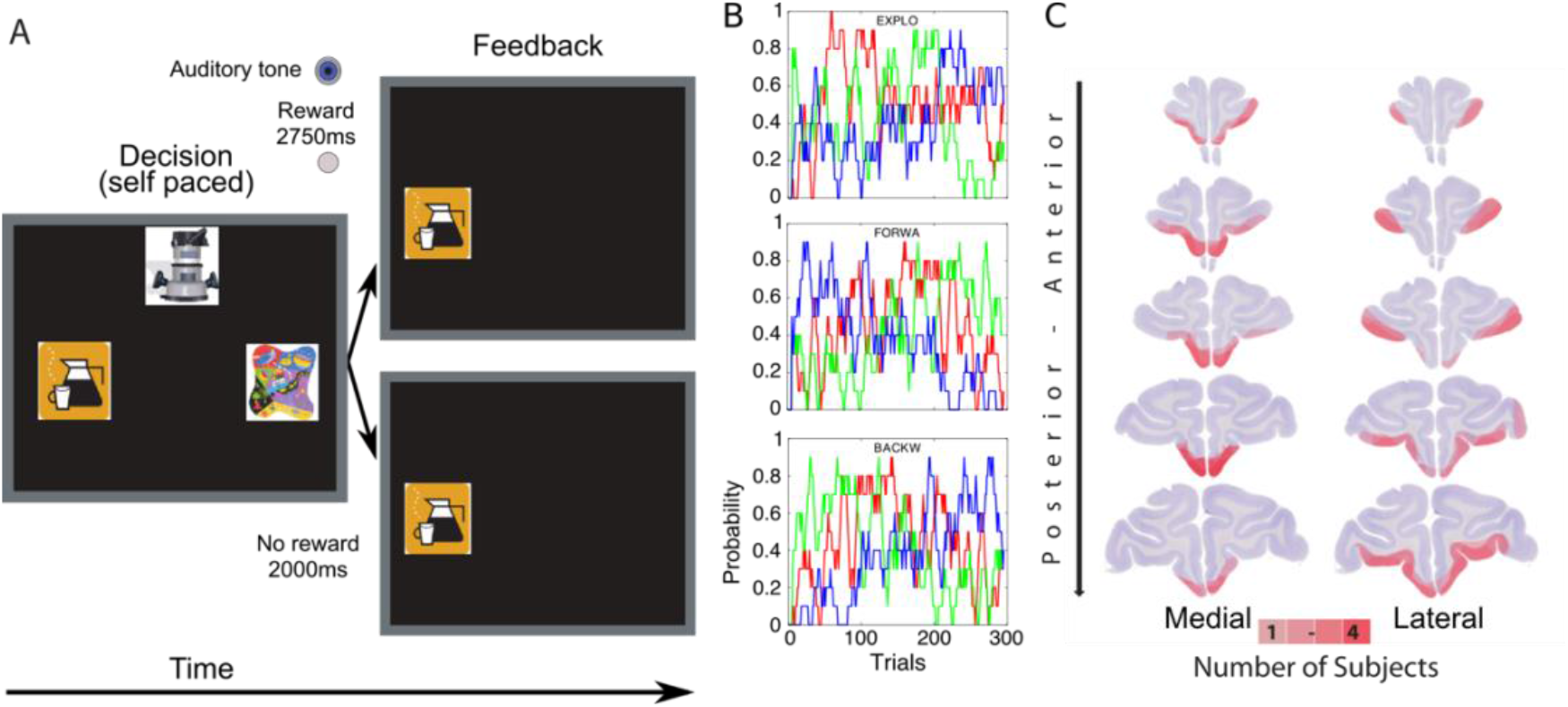
Task design, reward schedule and lesion. **(A)** Trial time line: Each testing session macaques made choices among three novel stimuli (novel clip art images; left-hand side) before receiving auditory feedback and, according to the particular reward schedule, a sucrose pellet reward (right-hand side). The chosen stimuli remained onscreen during feedback. Both possible outcomes are displayed in this example: A reward pellet was delivered in the case of a reward outcome and nothing happened during no reward events. **(B)** Animals made choices with three reward schedules in which the reward probabilities ranged between 0 and 1 and drifted throughout the session, with each option being competitive at some time during the session (i.e. each option was the best one at least during a short phase of the session). **(C)** Medial (Left) and lateral prefrontal (right) lesion locations represented on an unoperated control, with redness indicating lesion overlap (n=4 and n=3 respectively). Reward Schedules termed; Forwards (FORWA), Backwards (BACKWA), Exploration (EXPLO).

We used an established general linear model (GLM; see Methods) approach that analyses the tendency of animals to stay with or switch away from an option chosen on a given trial t (Wittmann *et al*., 2020). In this “credit assignment GLM”, we simultaneously account for several factors driving choice (Fig.2A). We started our analyses by comparing the effects of the contingent reward history (CxR-history) on learning between a baseline condition and the lateral lesioned animals using linear mixed effects models (LME) to account for monkey identity (Fig.2B,C). Typically, the extended contingent reward history incentivizes a learner to stay with their current choice (positive weights of CxR-history) and we would expect this effect to be significantly reduced in these lesioned animals if these regions are important for assigning rewards to the choices they are contingent on. Although, we did not find such a significant reduction in the most recent weight of CxR-history (RxC_t_: estimate-Lateral=0.004, SE=0.006; χ2(1)=0.391, p=0.532), inspection of subsequent time points revealed significantly reduced influence of contingent rewards on staying with a choice for RxC_t-1_ (estimate-Lateral=-0.016, SE=0.004; χ2(1)=7.986, p=0.005), RxC_t-2_ (estimate-Lateral=-0.020, SE=0.004; χ2(1)=8.330, p=0.004), and RxC_t-3_ (estimate-Lateral=-0.012, SE=0.003; χ2(1)=6.201, p=0.013). They align well with previous reports of a reduced ability to associate rewards with appropriate choices after lateral prefrontal lesions (Walton *et al*., 2010; Rudebeck *et al*., 2017). The fact that our new analysis approach picked these deficits up lays a solid foundation for the subsequent analysis of developmental changes of these effects in humans and motivates the hypothesis that contingent credit assignment is strengthened over the course of adolescence as lateral frontal cortex continues to mature.

**Figure2.**
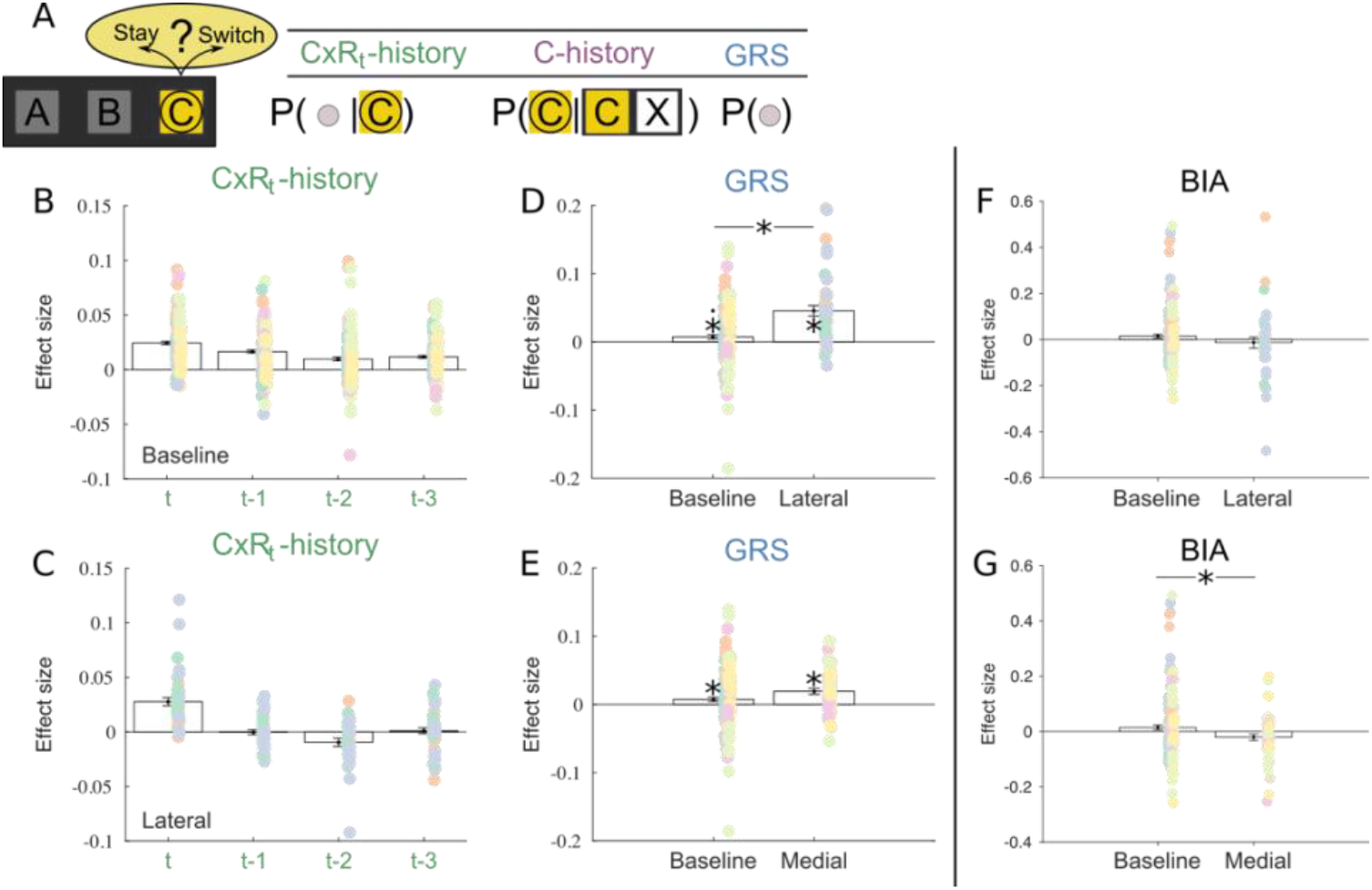
Credit assignment behaviour in macaques and the effects of lateral prefrontal cortex lesions. **(A)** Using a general linear model, we analysed macaque’s choices to stay with, or abandon, a currently pursued choice C. We analysed this decision as a function of previous contingent rewards for C (CxR_t_-history) as well as the global reward state (GRS) and also the pure choice history (C-history) in our “credit assignment GLM”. **(B**,**C)** Results of logistic GLM model predicting the tendency of macaque monkeys to stay with a current choice. Contingent credit assignment effects, the effects of CxR_t_-history on the decision to stay are significantly reduced after Lateral lesions (C) compared to a baseline control group (B). **(D**,**E)** In the same GLM model, the global reward state (GRS) exerts a positive effect on stay decisions; the animals are more likely to stay with a currently pursued choice if the average levels of rewards in the task are currently high. By including the effect in the same GLM as CxR_t_-history, the positive GRS effects account for variance in the choice data that is not explained by the contingent reward effects assigned by CxR_t_-history. GRS effects are significantly positive in the baseline group (D) and also in the Medial lesioned group (E). However, GRS effects are significantly increased after Lateral lesions. (D). **(F**,**G)** Finally, Lateral lesions do not affect biases induced by irrelevant alternatives (BIA) on choice (F), whereas Medial lesions do significantly decrease the bias by irrelevant alternative (BIA) effect compared to baseline (G). (dots indicate individual sessions; dot color indicates monkey identity; plots show mean -/+SEM, *, p<0.05)

After a consideration of contingent learning, we examined the influence of the GRS on staying with a currently pursued choice (Wittmann *et al*., 2020). Importantly, we analyzed these GRS effects in the same GLM model reported above assuring that any identified GRS effects could not be attributed to contingent reward learning. In this context, any GRS effect means that irrespective of whether the specific choice was or was not rewarded in the past, the overall average levels of rewards influence the animal in their decision to stick with a choice. We found such a significantly positive non-contingent effect of GRS in the baseline data (Fig.2D intercept-estimate=0.007, SE=0.003; χ2(1)=5.122, p=0.024) as well as in lateral lesioned macaques (intercept-estimate=0.046, SE=0.008; χ2(1)=8.210, p=0.004). Crucially though, the non-contingent effect of the GRS was changed in lateral lesioned animals compared to the baseline condition (Fig.2D, estimate-lateral=0.038, SE=0.008; χ2(1)=6.492, p=0.011). Neuroimaging work in macaques have identified neural correlates of the GRS in the agranular insular cortex (Wittmann *et al*., 2020); regions whose activity or connections might well have been impacted by the lateral lesions. This suggests that both contingent and non-contingent learning mechanisms rely on reward computations performed in and adjacent to lateral prefrontal cortex and motivates the hypotheses that both learning mechanisms may undergo developmental changes in humans over the course of adolescence.

We assessed the specificity of our lesion effects on GRS-based learning by comparing them to animals with lesions located in more medial parts of orbitofrontal cortex (referred to as *Medial* lesions). Like the baseline data and Lateral animals, Medial lesioned macaques also exhibited a positive GRS effect (Fig.2E, estimate-intercept=0.019, SE=0.004; χ2(1)=9.401, p=0.002). While the strength of this effect was not significantly stronger than observed in the baseline condition (estimate-Medial=0.012, SE=0.006; χ2(1)=2.517, p=0.113), it was significantly weaker than the one we observed in our Lateral condition (estimate-lateral=0.027, SE= 0.009; χ2(1)=5.080,p=0.024). In summary, this suggests that the influence of the GRS on credit assignment is particularly mediated by lateral regions of prefrontal cortex and these two subcomponents of reward learning thus represent likely candidate processes that might also continue to mature during human adolescence.

Finally, we contrasted these credit assignment mechanisms with computations that, in the macaque, do not rely on lateral prefrontal cortex but instead are linked to more medial parts of prefrontal cortex (Boorman *et al*., 2009; Basten *et al*., 2010; Papageorgiou *et al*., 2017). Medial frontal cortex carries decision signals in both humans and macaques, with lesions in this area affecting how the comparison of two options is influenced by an irrelevant third alternative (Noonan *et al*., 2010; Noonan *et al*., 2017). Following our analysis approach established in human medial orbitofrontal lesion patients, (Noonan et al., 2017), we used a combination of multinomial logistic regression analysis and reinforcement learning modelling (Fig.S1). We considered each three-choice decision as two binary comparisons and rearranged them such that we can extract the biasing effect of the value of a distractor option on choice (see Methods). We refer to this distractor effect as *bias by irrelevant alternative* (BIA) effect. In humans, BIA is significantly decreased after lesions to medial frontal cortex meaning that a relatively high value of the irrelevant alternative decreases the effect of the relevant value comparison on choice. Using the same analysis on the macaque data we replicate this pattern in the present analysis. Medial lesioned animals have a significantly decreased BIA effect compared to baseline (Fig.2G; Estimate-Medial=-0.054, SE=0.026, χ2(1)=1, p=0.048). By contrast, BIA in Lateral lesioned animals is not different from non-lesion control group (Fig.2F; Estimate-lateral=-0.027, SE=0.037, χ2(1)=1, p=0.533). This suggests that the effects of irrelevant alternatives on decision-making are mediated by medial parts of prefrontal cortex. As such, we might not expect this computation to mature in the same way during adolescence as those that rely on lateral regions as previous maps indicate the potentially protracted maturation of lateral orbitofrontal cortex (Gogtay *et al*., 2004; Raznahan *et al*., 2011). However, detailed comparisons of the neurodevelopmental trajectories of these regions are still missing to our knowledge. We will test the developmental trajectories of these reward and decision-related computations empirically in the following sections applying the same analyses techniques as those used in this section to human data.

### Probabilistic reward learning performance increases during human adolescence

We have shown that, in macaques, lateral regions of the prefrontal cortex are linked to two important ways to assign rewards to choices: contingent credit assignment, and non-contingent effects of the GRS. Next, we tested the hypothesis that these credit assignment computations continue to change during human adolescence when parts of particularly lateral prefrontal cortex continue to mature *(Gogtay et al., 2004; Raznahan et al., 2011)*. We collected data from human participants on a matched 3-choice probabilistic decision-making task akin to that used in the macaque study (Fig.3) and employ the same first-level analyses to assess the influence of contingent rewards and the GRS on learning. Again, we contrast the computations with computations linked more specifically to medial parts of frontal cortex, in particular the *biasing effect of irrelevant alternatives (BIA)*.

**Figure 3.**
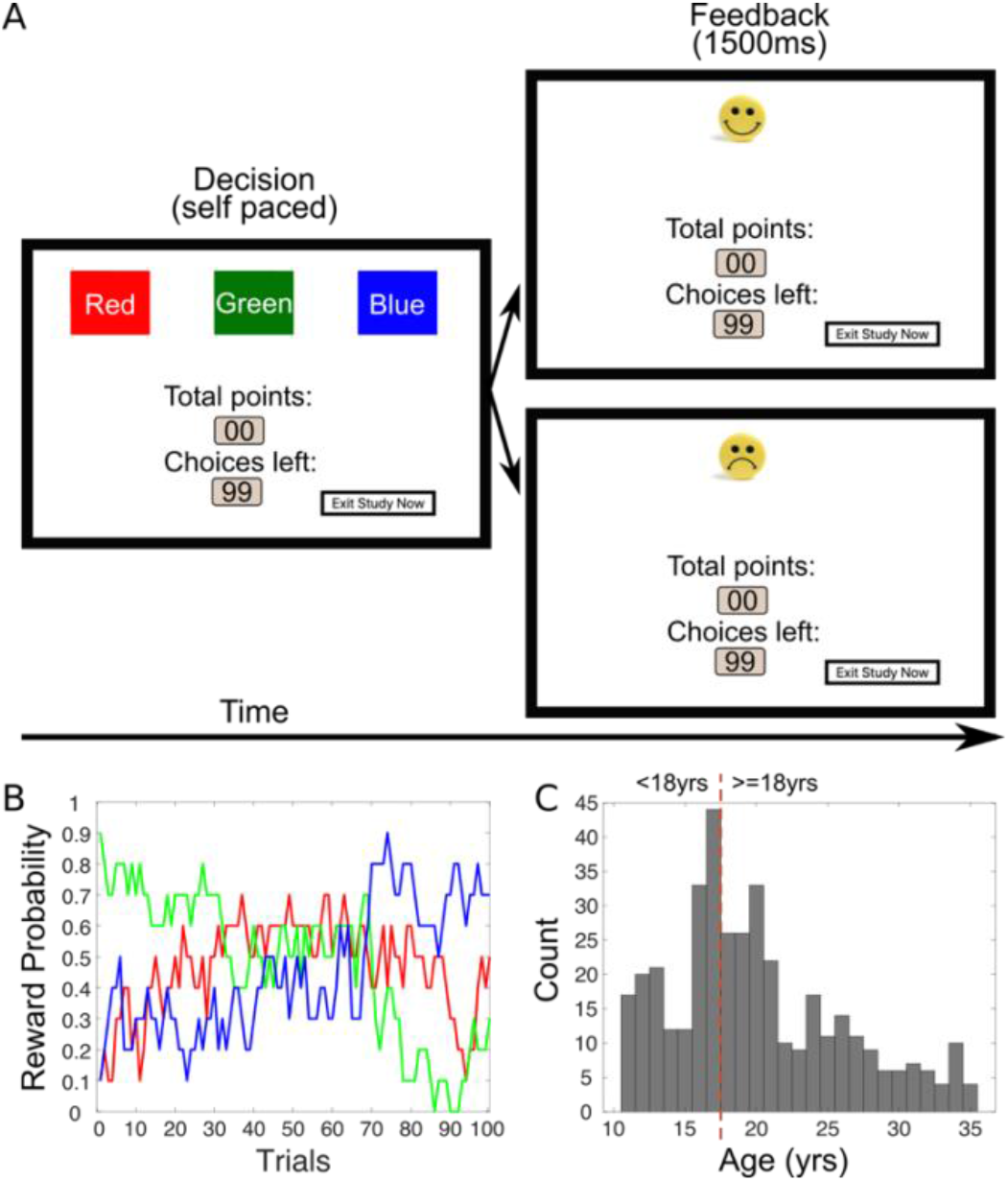
Task design, reward schedule and sample. **(A)** Trial timeline: Participants decided between three choice options (red, green, blue squares; left-hand side) before receiving feedback for 1500ms (right-hand side) whether their choice yielded a reward (10 point and smiley face) or no reward (no points and sad face). Both possible outcomes are displayed in this example. **(B)** Reward probabilities ranged between 0 and .9 and drifted throughout the session. **(C)** Age distribution of the final sample with dashed line indexing the age groupings cut off at 18 years. We refer to participants younger than that age as *adolescents* and we refer to older participants as *young adults*.

Before progressing to our key analyses, we first assessed broad measures of task performance in our human experiment and their change from adolescence to young adulthood. We found that overall task performance as measured by total rewards acquired in the experiment increased across age. Young adults earned more total rewards than adolescents (t_386_=3.47, p=0.001) and there was a linear correlation between total rewards and age across our age range of 11 to 35 years (Fig.4A; R=0.16, p<0.001). Follow-up model fits suggest that the relationship of age with total rewards was best characterised by a quadric function (R=0.16, p<0.001, LRT: p=0.0039, R^2^=0.03). In accord with better overall performance, the frequency with which the highest value-option (as defined by value estimates from a reinforcement learning model) was chosen was significantly higher in young adults compared to adolescents (Fig.4B; t_386_=7.89, p<0.001). Again, this was a linear increase with age (R=0.29, p<0.001, LRT: p<0.001, R^2^=0.11).

**Figure 4.**
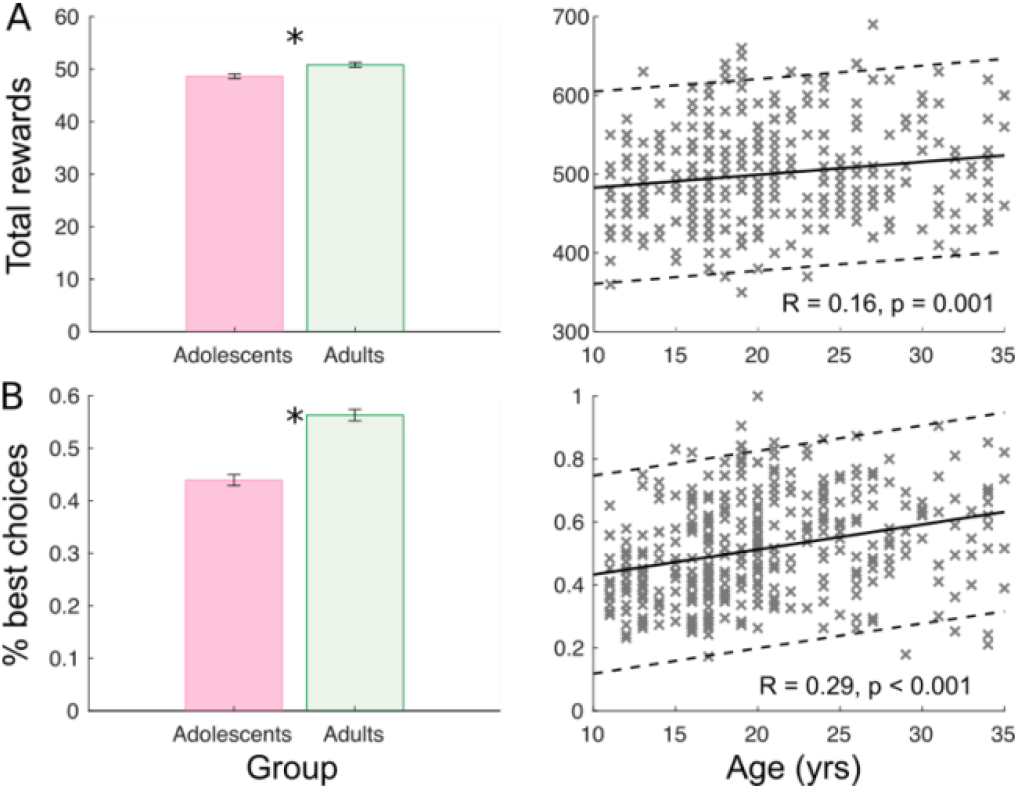
Performance in probabilistic 3-choice learning task increases during adolescents. **(A)** Young adults, compared to adolescents, earned more total rewards in the experiment (left) and there was a linear increase in the total rewards earned across age (right). **(B)** In line with this, the frequency of choosing the highest-value option was higher in young adults (left) and also increased across age (right). Note that chance performance would be at 0.33 since it is a three-choice task. (“x”s indicate individual participants; left plots show mean -/+SEM; solid line in the right plots indicate best fitting linear trend, Dashed line represented 95^th^ % confidence interval; * p<0.05)

### Contingent and GRS-based credit assignment change during human adolescence

We went on to apply our key analysis, the *credit assignment GLM*, to discern whether these changes in task performance coincide with the maturation of dissociable subcomponents of reward learning – contingent credit assignment (CxR-history effects) and learning from the GRS. Such changes are predicted from our macaque results suggesting that both processes rely on lateral regions of prefrontal cortex which matures very late in humans.

First, we examined the effects of CxR-history. Whether the most recent choice was rewarded or not should be the largest predictor of staying with this choice. As a whole sample (regardless of age) participants stayed with an option if they had been rewarded on the current trial (t) for choosing it (one-sample t-test; t_352_=10.92, p<0.001). Indeed, comparing the effect sizes of CxR_t_ between adolescents and young adults showed that the size of this effect was bigger in young adults (t_351_=4.34, p<0.001) suggesting increasing associability between rewards and appropriate choices. Furthermore, correlating CxR_t_ with age confirms this as a linear relationship (Fig.5A; R=0.22, p<0.001, LRT: p<0.001, adjusted R^2^=0.050). By contrast, we found no developmental changes in reward-unrelated C-history effects (Fig.S2).

**Figure 5.**
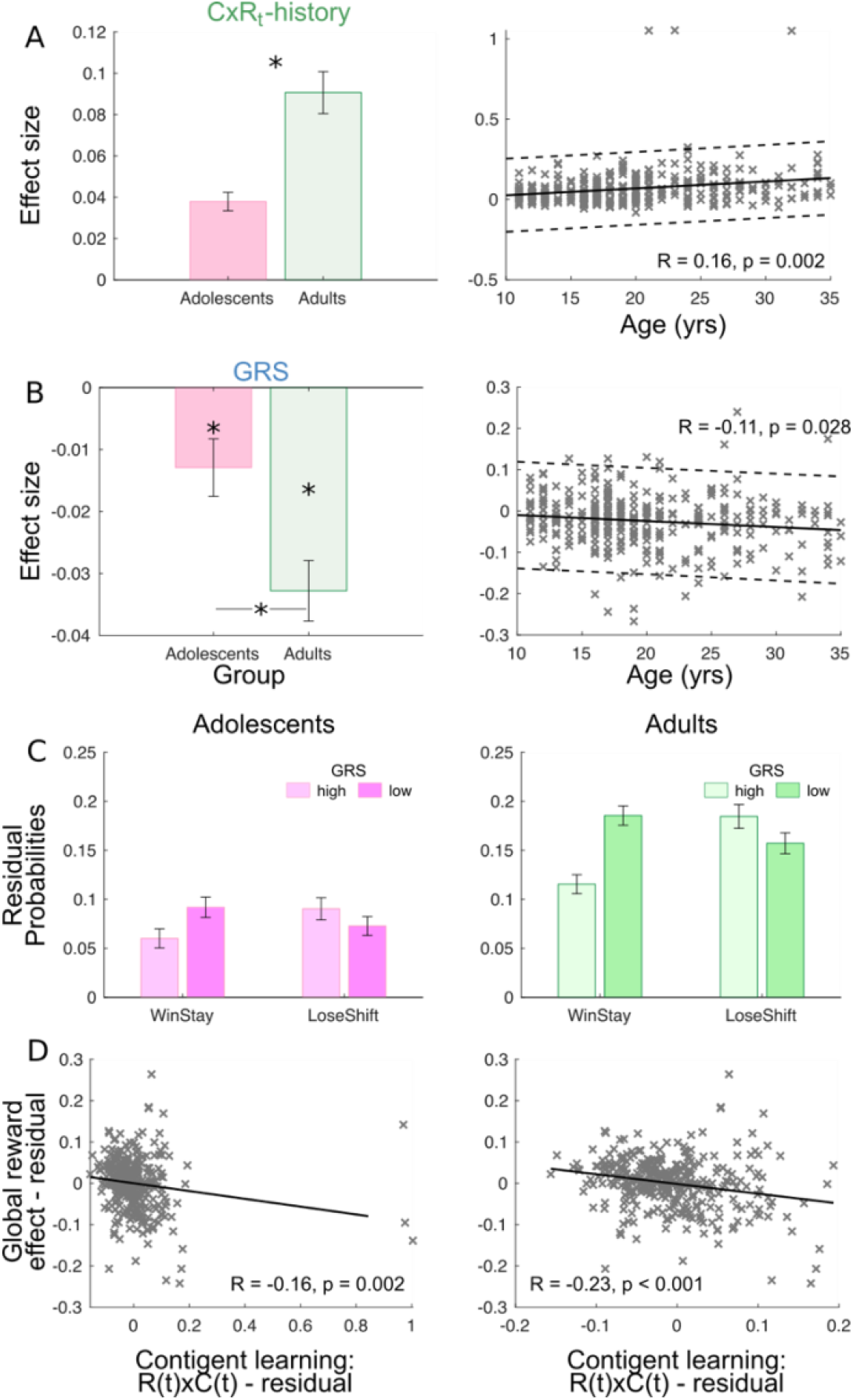
Contingent credit assignment increases across age and negative GRS effects decrease across age. We applied the same *credit assignment GLM* we used in the macaque experiment to analyze the developmental trajectory of credit assignment computations during human adolescence. Panels A and B show effect sizes for component parts of this GLM. **(A)** Considering the effect of the most recent outcome on the tendency to repeat a choice (CxR_t_ effect), we found that young adults had a stronger tendency to repeat rewarded choices compared to adolescents (left), with effect size linearly increasing with age (right). **(B)** Independent and in addition to contingent credit assignment, GRS had a negative effect on staying with an option. Participants tended to stick more with a choice if it was encountered in the context of a low overall GRS. Such negative GRS effects were found in both adolescents and young adults with a significant difference between them (left). This indicates that young adults had an even stronger tendency to contextualise rewards by the global reward state and this is replicated by a linear decrease of GRS over time (right). **(C)** Plots show residual probability of staying after a win and shifting after a loss separated by low and high GRS (light and dark grey, respectively) for adolescents (left) and young adults (right). Note that an overall increase in residual probabilities of choosing to stay after a reward (WinStay) or switch after a loss (LoseSwitch) both increasing from left to right panels from adolescents to adults. As this effect of contingent learning increased, also the effect of the GRS as indexed by the interaction of GRS with WinStay/LoseShift increased. As people get older, they were even more likely to repeat rewarded choices when encountered in a low GRS (darker bars) and simultaneously are more likely to switch away from losing choices if encountered in a high GRS (lighter bars). **(D)** Finally we examined the relationship of GRS with contingent learning independent of age. We conducted this as a partial correlation after accounting for possible confound factors. This analysis revealed a strong negative correlation suggesting that individuals whose choices are guided by contingent learning mechanisms are also likely to be negatively influenced by the GRS. (“x”s indicate individual participants; plots show mean - /+SEM; solid line in the right plots indicate best fitting linear trend, Black line represents linear best fit. Dashed line represented 95^th^ % confidence interval. * p<0.05).

Then, we examined the effects of the GRS in our GLM. We have shown that GRS effects are positive in macaque monkeys and we have shown that these effects were even increased after Lateral lesions. We now consider GRS effects in our human sample. Again, we find effects of GRS on the decision to stay with a choice. However, strikingly, in our sample as a whole, we did not find a positive, as in the macaques, but instead a significantly *negative* effect of GRS (t_352_=6.98, p<0.001). The effects were significantly negative in both the adolescent (one-sample t-test t_154_=-2.78, p=0.006) and the young adult group (t_197_=-6.72, p<0.001). That means that irrespective of directly reinforced choices, if participants had observed many rewards in the past (high GRS) then they were more likely to switch away from a current choice. By contrast, if GRS was low, indicating the absence of better alternatives in the past, then participants continued pursuing their choice even in the absence of contingent reward. Importantly, as predicted from the animal result, the GRS effect changed during adolescence. The GRS effect was even more negative in young adults compared to adolescents (Fig.5B; t_351_=-2.89; p=0.004) and correlated linearly negatively with age (R=-0.14, p=0.011; LRT p=0.0379, R^2^=0.018).

These results suggest that the GRS alters the behavioural response to rewards received for a current choice. To examine the effects of the GRS more directly we followed an approach used by Wittmann and colleagues (2020) and regressed all effects of the previous GLM, with the exception of CxR_t_ and GRS, out of the stay/switch decision data (Fig.5C). We then binned the residual choice probabilities by the receipt of a reward on trial t (i.e. the CxR_t_ effect), a median GRS split and by age (adolescents and adults <18/≥18). Based on whether CxR_t_ was rewarded or not and the consequent stay or shift response, we coded the residual probabilities in terms of winStay and loseShift and this allowed us to directly examine how the GRS impacted winStay/loseShift behaviour. This analysis shows an interaction of winStay/loseShift and GRS independent of age group (F_1,385_=79.15, p<0.001) illustrating the GRS effect observed before: while participants were more likely to stay after a reward, they did this even more in a low GRS; in a high GRS, they were quicker to switch away from unrewarded choices. However, critically, in addition to a main effect of age (F_1,384_=36.48, p<0.001), the GRSxWinStay/loseShift interaction changed with age group in a manner suggesting that adolescents were relatively less influenced by the GRS in value updating (WS/LS x GRS x age: F_1,384_=7.27, p=0.007). Older participants, by contrast, showed a stronger contrast effect after receiving reward: In low GRS environments they were particularly likely to stay with rewarded options and less likely to switch away from unrewarded ones.

Our findings show that both contingent credit assignment and non-contingent assignment of the GRS to choices changes during adolescence. These effects are predicted from the same analysis pipeline applied to macaque data on the same task, and the fact that lateral orbitofrontal cortex matures particularly late during human development. Strikingly, the GRS effects are reversed in humans compared to the macaques suggesting a more optimal use of the GRS to “contrast” and contextualize current reward effects (Crespi, 1942; Daw & Touretzky, 2002; McNamara *et al*., 2013).

### Contingent credit assignment and negative GRS effects correlate across participants

As a last step in our examination of contingent and non-contingent credit assignment, we investigated the relationships between contingent reward assignments and a negative GRS effect. Despite the theoretical accounts cited above arguing that a negative GRS effect might aid adaptive value learning, it might be argued that GRS effects per se are suboptimal in the context of probabilistic learning tasks. To address this empirically, we examined the relationship of GRS with a marker of contingent learning, the CxR_t_ effect, as the latter reflects a signature of adaptive learning in the context of this task. Controlling for participant age and their GLM constant, we correlated GRS and RxC_t_ using a partial correlation. Our findings revealed a strong negative correlation between contingent reward assignment and the global reward effect (Fig.5D right; R=-0.16, p=0.002). This suggests that individuals who are more influenced by contingent reward assignment mechanisms are also more likely to rely on a negative reward contextualisation. This pattern of behaviour further supports the idea that negative GRS effects are adaptive and may co-mature with contingent credit assignment mechanisms during adolescence. Visual inspection might suggest that the correlation is potentially driven by three outliers with high contingent learning scores. However, removal of these data points confirmed that this was not the case; instead, the correlation became even more significant (Fig.5D left; R=-0.23, p<0.001).

### No developmental changes in computations linked to medial parts of prefrontal cortex

Having established developmental changes in contingent and non-contingent credit assignment – both linked to lateral prefrontal cortex – we asked whether decision computations supported by closely adjacent parts of medial frontal cortex may or may not exhibit a similar developmental change during adolescence. As in our macaque analyses, we first fitted a simple reinforcement learning model to our data. In line with the increased effects of CxR_t_ reported above, we found that learning rates for young adults were significantly higher than for adolescents (Fig.6A; t_386_=-3.83, p<0.001) and again this result was confirmed by a linear relationship across age (R=0.19, p<0.001, LRT: p=0.001, R^2^=0.035). Notably, the age groups did not differ in their general levels of decision-making noise, as the RL models’ inverse temperature parameter did not differ with age (Fig.6B; t_379_=-1.83, p=0.068; R=0.02, p=0.720). Note the effect is still non-significant when the inverse temperature is not log normalized. This suggests that changes in learning rates cannot be reduced to changes in decision noise. However, it might be that more sensitive analyses and larger sample sizes can uncover co-occurring changes in the inverse temperatures.

Based on the RL model, we repeated our multinomial logistic regression to isolate the *bias by irrelevant alternative* (BIA) effect (see Methods). If the medial parts of frontal cortex linked to the BIA effect are developmentally delayed, like the lateral parts, then we would hypothesize a negative BIA effect in younger participants and would abate with age. A negative effect would mean that, despite the third option being, in principle, irrelevant, when its value in each binary comparison is high (versus low), decisions between the relevant options are less influenced by their value difference, a pattern seen in decision strategies such as divisive normalization mediated by regions such as the parietal cortex (Louie *et al*., 2011; Louie *et al*., 2013; Chau *et al*., 2014). However, this pattern was not evident in the data. The overall positive effect of the BIA (one-sample t-test; t_340_=3.25, p=0.001) did not show between-group differences or correlate with age (Fig 6C; t_339_=1.06, p=0.291; LRT: R=0.03, p=0.522). Therefore, in contrast to the credit assignment mechanisms linked to lateral prefrontal cortex, the influence of the value of third option on the binary choice was stable across the ages we tested.

**Figure 6.**
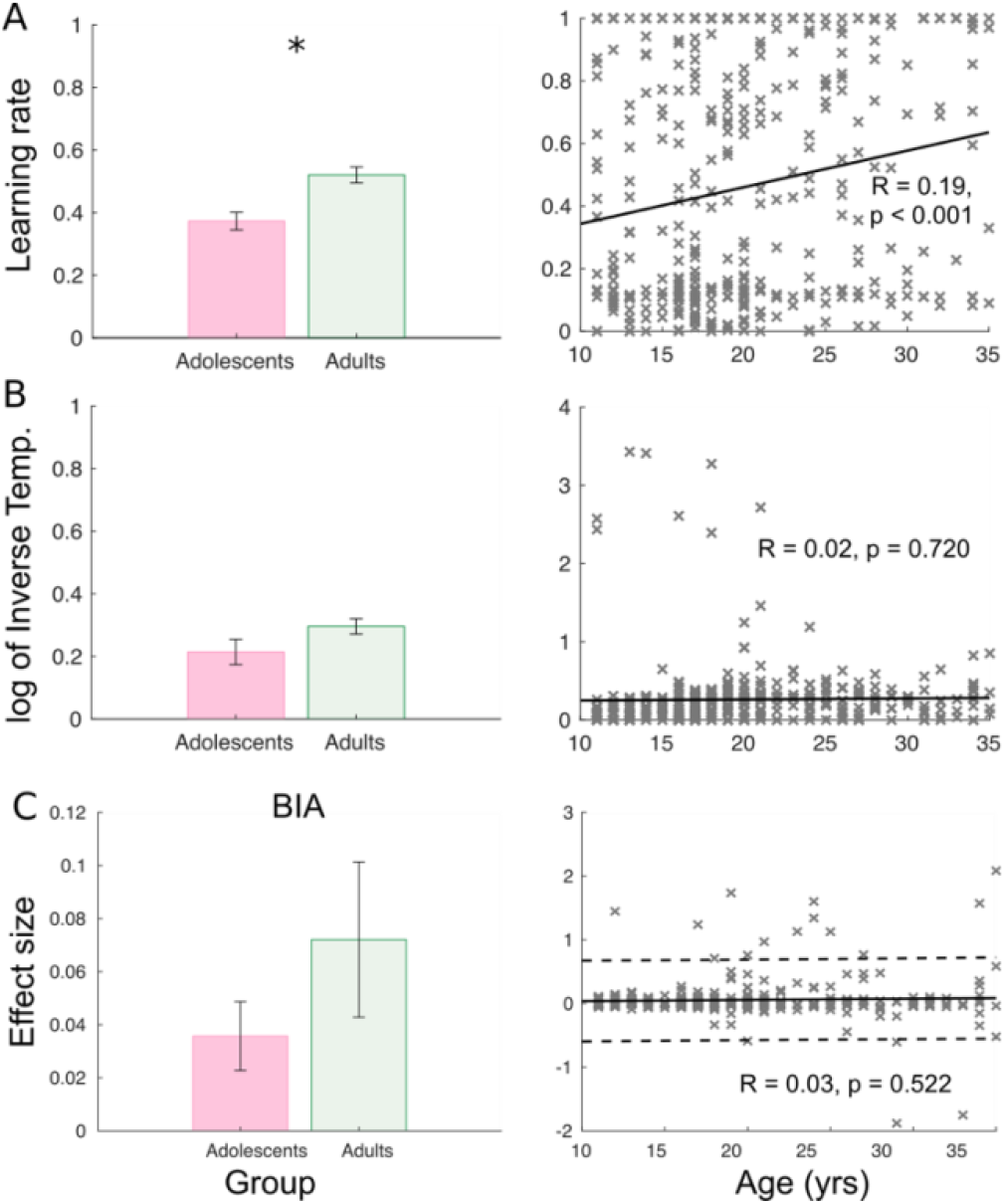
Reinforcement model parameter fits and influence of the irrelevant alternatives on choice. **(A)** The learning rate increased in young adults compared to adolescence (left) and increased linear (right). **(B)** By contrast, the log of the inverse temperature does not show a significant difference in the two age groups (left), nor a significant increase across age (right). **(C)** BIA effects on choice do not change with age. (“x”s indicate individual participants; plots show mean -/+SEM; solid line in the right plots indicate best fitting linear trend, Black line represents linear best fit. Dashed line represented 95^th^ % confidence interval. * p<0.05).

## Discussion

Adolescence is a phase of wide-ranging changes in cognitive functioning and brain development (Gogtay *et al*., 2004; Shaw *et al*., 2006; Raznahan *et al*., 2010; Dahl *et al*., 2018). We have shown that during this phase two important component processes of reward learning continue to mature: contingent credit assignment and non-contingent learning based on the GRS. The GRS reflects the average reward levels available in an environment independent of the specific choices made. These learning mechanisms have been linked to specific parts of lateral prefrontal cortex in animal studies (Walton *et al*., 2010; Rudebeck *et al*., 2017; Wittmann *et al*., 2020). Using finely matched experimental paradigms and analysis techniques, we demonstrate that both the effects of contingent rewards and the GRS on learning are causally impacted by lateral prefrontal lesions in macaque monkeys. Together with the observation that this part of the human brain undergoes a particularly protracted maturation process (Gogtay *et al*., 2004; Raznahan *et al*., 2011), our results support the idea that neural maturation of this part of the brain drives the maturation of both contingent and GRS-based learning during adolescence in humans. Our species comparison further reveals a qualitative difference in GRS effects. While, in macaques, the GRS appears to cause noise in the learning process by assigning the credit of preceding rewards to multiple outcomes, our human participants appear to use the GRS in a qualitatively different manner to contrast new rewards with the baseline level of rewards encountered in the past – a process that can support optimal choice switching and exploration (Daw & Touretzky, 2002; McNamara *et al*., 2013). Consistent with an adaptive role of the GRS in human credit assignment, we found that – independent of age – participants that are strongly influenced by the GRS are also able to assign more credit to contingent rewards.

The ability to form specific links between stimuli and outcomes required for contingent learning has been strongly linked to orbitofrontal cortex (Noonan *et al*., 2017; Howard & Kahnt, 2018; Groman *et al*., 2019; Knudsen & Wallis, 2020). We have replicated such deficit in contingent reward assignments using the same analysis pipeline we applied to our human data on a large data set of macaque data including macaques with aspiration lesions in lateral prefrontal cortex (Fig.1) (Noonan *et al*., 2010; Walton *et al*., 2010). Beyond replicating previous results, we have demonstrated that the same lesions impact not only contingent credit assignment, but also learning based on the GRS (Fig.2D). This means that, after lateral prefrontal lesions, not only are individual rewards less strongly assigned to the specific choices that caused them, but they also spread more broadly to unrelated choices (Thorndike, 1933b) suggesting a general increase in the imprecision of reward credit assignments after lateral prefrontal lesion. However, it is important to consider the anatomical specificity of the lesion effect in more detail. Although our lesions targeted Walker areas 11 and 13 in lateral orbitofrontal cortex, recent work indicates that aspiration lesions to this part of the brain also disconnect adjacent posterior lateral prefrontal cortex (Rudebeck *et al*., 2013). While our observed deficit in contingent learning is likely to be caused by damage to the orbital part of area 12 (Rudebeck *et al*., 2017; Murray & Rudebeck, 2018), there are reasons to think that the GRS effects are mediated by more posterior parts of lateral prefrontal cortex that are disconnected by the lesion, in particular anterior agranular insular cortex bordering orbitofrontal cortex. First, a recent study suggests that GRS effects on choice rely on a representation of the GRS in macaque agranular insular cortex (Wittmann *et al*., 2020) and also human anterior insula carries similar reward signals (Wittmann *et al*., 2016). Second, the paradigm employed by the latter animal study and our study are variants of the classic reversal learning paradigm and for this type of task, it is bilateral agranular insula that undergoes most profound grey matter during training (Sallet *et al*., 2020). Together, this suggest that adjacent parts of lateral prefrontal cortex carry choice-specific as well as global reward signals important for guiding learning in reversal learning type tasks.

Using the analogous analysis pipeline and three-choice probabilistic task, we have demonstrated that the GRS impacts learning in monkeys and humans. We have shown that lateral prefrontal lesions alter macaque GRS effects and that, concordantly, the influence of the GRS on learning increases during adolescence in humans. In all analysed macaque data sets here and consistent with previous work on the GRS (Wittmann *et al*., 2020) and credit assignment in general (Walton *et al*., 2010; Rudebeck *et al*., 2017), the effect of the GRS on choice was *positive*. Positive GRS effects indicate that choices made in high GRSs are more likely to be repeated, even if the current choice results in failure. In sharp contrast, in humans, the GRS exerted a *negative* effect on staying with a choice. This indicates that when the GRS is very high, people are more likely to switch away from a choice that is not rewarded while when the GRS is low, they are more likely to continue pursuing unrewarded options. Such negative contrast effects of the GRS can subserve optimal switching in natural environments because they can incentivize persistence when the reward environment is sparse, but encourage exploitative strategies in high reward environments (Crespi, 1942; Daw & Touretzky, 2002; McNamara *et al*., 2013). These considerations suggest that the GRS can guide learning in macaques and humans in qualitatively different ways during probabilistic learning tasks.

The representations adolescents held of the GRS impacted negatively on their decisions to stay with a choice or to switch to an alternative. This tendency was strengthened towards young adulthood, reflecting an even stronger referencing of current outcomes to the GRS (Crespi, 1942; Daw & Touretzky, 2002; McNamara *et al*., 2013). This observation may provide an avenue towards understanding changes in adolescent attitudes towards exploration and risk and uncertainty from a mechanistic perspective (Hartley & Somerville, 2015; van Duijvenvoorde *et al*., 2016; van den Bos & Hertwig, 2017). For instance, our results indicate that adolescents may display stronger persistence with unrewarded options, for example when the GRS is high. In such a setting, young adults by contrast more readily refrain from exploration and switch away from an unrewarded choice as the high GRS discourages exploring new choice options and incentivizes switching back to previously rewarded options. The fact that negative GRS effects are correlated with contingent credit assignment (Fig.5D) might further suggest that both mechanisms are part of general developmental trajectory towards more exploitative task strategies. Such increased exploitative strategies might be evidenced by the increase in task performance in general and specifically by the concurrent increase in contingent learning (and correspondingly the learning rate from our reinforcement learning analysis) across age. This is in line with age-related differences of the impact that outcomes have on value updates between these age groups have been reported before (van den Bos *et al*., 2012; Hauser *et al*., 2015; Xia *et al*., 2020). Rather than reflecting more optimal learning per se (Davidow *et al*., 2016; Lloyd *et al*., 2020; Lockwood *et al*., 2020), the benefits of negative GRS effects, just as the ones of increased contingent learning, might be felt particularly in environments where exploration is relatively discouraged and choices should be directed towards options with high values and at the expense of sampling more uncertain options that nonetheless might prove more beneficial in the long run (Daw *et al*., 2006; Wilson *et al*., 2014; Trudel *et al*., 2020). Our results may suggest that part of the developmental changes underlying these behaviors are rooted in a specific subcomponent of reward learning that rely on the GRS, and that is linked to late maturing lateral parts of prefrontal cortex. Therefore, an increased focus on the contribution of the prefrontal cortex in addition to subcortical structures important for value coding (Galvan & McGlennen, 2013; Mills *et al*., 2014; Insel *et al*., 2017; Vaghi *et al*., 2020) might in the future improve our understanding of adolescent risk taking, learning and choice.

One final contribution of this study comes from the evidence of heterogeneity across the developmental profiles of the three learning and decision-making processes. Unlike the reward learning mechanisms, we find no evidence of age-related differences with respect to the influence on choices of high values of irrelevant alternatives, in the age range we tested. If all orbitofrontal subregions mature at the same rate then we would predict that the decision-making mechanism supported by the medial orbitofrontal cortex would also correlate with age we would have observed choices that violated the axiom of irrelevant alternatives. Contrary to these hypotheses the degree to which a high value irrelevant option influenced choice did not differ across age. If adolescents are already able to use adult-like decision-making mechanisms it may suggest that the medial orbitofrontal cortex may have already matured by this age. These findings raise the possibility of functionally meaningful fine-grain differences in the rate of structural maturation between these orbitofrontal subregions which to date, while visually indicated by current whole brain developmental maps (Gogtay *et al*., 2004), has not been anatomically quantified.

## Methods

### Subjects: Macaques

Data from six male rhesus macaque monkeys (Macaca mulatta), aged between 4 and 10 years and weighing between 7 and 13.5 kg was reanalysed for the purposes of the current experiment. This data was originally collected and analysed in Walton et al (2010) and Noonan et al (2010). Six monkeys originally participated in an experiment reported by Walton et al (2010) in which three animals acted as unoperated controls, whereas the other three received bilateral aspiration lateral orbitofrontal cortex lesions following training and presurgical testing. The three unoperated control monkeys, and one additional monkey who had not participated in Walton et al study then participated in Noonan et al (2010) and were tested before and after bilateral aspiration lesions of medial orbitofrontal cortex. All animals were maintained on a 12 hr light/dark cycle and had 24 hr ad libitum access to water, apart from when testing. All experiments were conducted in accordance with the United Kingdom Animals Scientific Procedures Act (1986).

### Subjects: Human adolescents and young adults

Participants between 11 and 35 years old were recruited. In total 422 participants completed the task. We refer to participants younger than that age as *adolescents* and we refer to older participants as *young adults*. Data was excluded from the analysis for failing to supply age or gender data. Participants were also excluded if they only repeatedly chose one option or one location, indicated they had completed the game more than once already or did not have parental permission. A further 7 participants were excluded from that sample as their median reaction times was three times the standard deviation from the mean. This left 388 participants (260 female, median age = 19).

Participants 18 years old and above were recruited and directed to the study website via University of Oxford email lists, public and university online direct advertisement and social media. Participants under 18 years old were recruited via their parents and schools within the Oxfordshire area. Collaborating schools forwarded documentation inviting parents to consent to their children participating in the study by directing them to the study website. Children or adults assented or consented respectively to participation in the study before the task began and had the option not to submit their data to the study once the task and questionnaires had been completed. The study was approved by the Central University Research Ethics Committee (Project Number: R59372/RE001). Participants were provided with no monetary compensation.

### Task and Procedure: Macaques

Apparatus, training histories and schedules have all been fully described in Walton et al (2010) and Noonan et al (2010). For the purposes of the present study, however, we will briefly describe the task and reward schedules (Fig.1A,B). On every testing session, animals were presented with three novel stimuli which appeared in one of four spatial configurations. Configuration and stimulus position was determined randomly on each trial. Stimuli remained on screen until an option was chosen. Reward was delivered stochastically for a choice towards each option according to the reward probabilities defined by the session schedules. Stimulus presentation, experimental contingencies, and reward delivery was controlled by custom-written software. Here, we analyse data from three reward schedules employed by these studies (Forwards FORW, Backwards BACKW, Exploration EXPLO) and which formed the basis of the experimental schedules used for our human study. In these schedules, all three options were at some point competitively rewarded and reward probabilities varied over the course of the testing session. The probabilities of each option being rewarded were independent of each other. Across different days, animals completed five sessions of 300 trials under each schedule, with novel stimuli each time. For FORW and BACKW, the sessions were interleaved across testing days, whereas for EXPLO the data was run with consecutive sessions. Data were collected both pre- and postoperatively. Approximately 18 months separated testing in the Walton et al. experiment and training in the Noonan et al study. Before testing in the latter study, all animals with medial orbitofrontal lesions were brought to a criterion of 80% correct on three choice-reversal schedules and both pre-operative groups were at roughly the same preoperative performance level as they were when they acted as unoperated controls in the former study.

Surgeries: Surgical procedures and histology for the lateral and medial prefrontal lesioned animals have been previously described in full in Rudebeck et al (2008) and Noonan et al (2010). In brief, animals were given aspiration lesions to the lateral or medial orbitofrontal cortex using a combination of electrocautery and suction under isoflurane general anesthesia. The lateral lesion was made by removing the cortex between the medial and lateral orbitofrontal sulci and as such predominantly targeted Walker’s areas 11 and 13, but may also have included parts of area 12. Medial lesions removed cortex between the medial orbitofrontal sulcus and the rostral sulcus; mainly including Walker’s area 14, but may have also included some parts of area 10. Note, the lateral aspiration lesion effects on contingency learning reported by Walton et al (2010) and Noonan et al (2010) have recently been argued to be caused not by cortical damage to Walkers area 11 or 13 but by the damaged cortex laying adjacently lateral to this area beyond the lateral orbitofrontal sulcus which transitions into ventrolateral prefrontal cortex and aligns mostly with the gyral region of the orbital part of inferior frontal gyrus (Rudebeck & Murray, 2014). This corresponds to the orbital part of area 12 (12o) in macaques and Brodmann’s area 47o in humans (referred to from here as area 47/12o). The contingency learning effects are now attributed to the disconnection of the lateral orbitofrontal cortex from other circuits via white matter damage to tracts traversing under central orbitofrontal cortex. We therefore refer to these lesions as “Lateral” and to the medial orbitofrontal lesions as “Medial”.

### Task and Procedure: Humans

Human participants completed a similar three-armed probabilistic bandit task (Fig.3A) that was in fact modelled after the monkey paradigms. It was coded in JavaScript, HTML and CSS and hosted on JATOS version 3.3.4. During the task, participants saw three different coloured options (blue, green and red), which were presented in one of three locations that varied along the x axis. As in the monkey study, the options locations were randomized across trials. By clicking on one of the options, participants would either see a smiley face and win 10 points or see a sad face and win no points. The goal was to win as many points as possible. At the bottom of the screen, throughout the game, the number of points participants had won so far and how many trials they had left to play was displayed. Also, participants were specifically instructed that the “chance of winning points is different for each color” and that throughout the game “the most rewarding color might change”. The reward schedule (Fig.3B) was adapted from one of the above macaque studies, Noonan et al. (2010). Reward probabilities were slowly and unpredictably drifting over time and ranged between 0 and .9 and resembled the monkey schedules. The probabilities of each option being rewarded were independent of each other. The task was self-paced, with stimuli remaining on the screen until a decision was made and feedback was presented for 1500 milliseconds. After a 10-trial practice run, participants could either go back to the instructions, if they had remaining questions, or proceed to the main task, which consisted of 100 trials and took approximately 7 minutes to complete. Participants also submitted age and gender information.

### First level analyses applied to both human and macaque data: Credit assignment general linear model (GLM)

For all behavioural analyses, we used MATLAB 2020Ra (The MathWorks) and SPSS (v25). We applied an identical first-level general linear model (GLM) to both human and macaque choice data (Fig.2A). Via the GLM, we sought to understand the factors which influenced the decision to stay or switch from a current choice in relation to contingent reward assignments, choice repetition irrespective of reward, and, importantly, the global reward state (GRS). The latter variable captures the average recent reward levels irrespective of the specific choices that have led to reward. In monkeys, a high GRS can increase the degree to which animals stay with their currently pursued choice even if this specific choice was not rewarded (Wittmann *et al*., 2020). We adapted the logistic GLM used in Wittmann et al (2020). For every trial t we identified the chosen stimulus C and examined whether it was chosen again on the next trial. We then tested whether such a stay/switch decision was predicted by three sets of regressors: 1. The contingent choice-reward history of C (CxR-history), 2. The reward-unlinked choice history of C (C-history) and, 3. the choice-unlinked reward history (GRS). The GRS regressor reflects our parameterization of the GRS and allowed us to test whether the global reward state (GRS), regardless of the choice history and the contingent choice-reward history, influenced stay/switch decisions. The regressors were constructed in the same way as in our past report (Wittmann *et al*., 2020) as follows:

1. <u>CxR-history:</u> The model captures contingent reward effects (CxR-history) through regressors that denote whether choices of C on trial t and also on the preceding three trials were rewarded or not (CxR_t_, CxR_t-1_, CxR_t-2_, CxR_t-3_,). CxR-history regressors were set to 1/0 for rewarded/unrewarded outcomes. Hence, positive effects of these variables indicate that a choice is more likely to be repeated if that specific choice has received reward in the past. Note t refers to instances in which choices of C are made and not necessarily to its presentation on consecutive trials as only the former inform the conjunctive choice-reward history of C.
2. <u>C-history:</u> The model includes three regressors to reflect the recent choice history of C (C-history; C_t-1_ C_t-2_, C_t-3_). Irrespective of the receipt of reward, this regressor codes whether C was chosen or not, being set to 1/0 for each trial. In contrast to CxR-history, C-history captures the degree of choice repetition, i.e. the fact that past choices predict that these same choices are made in the future independent of the receipt of reward.
3. <u>GRS:</u> The model took the simple average reward on the three trials before t an index of the overall current levels of reward. As with C-history (but unlike CxR-history), the three most recent trials were used for this in all cases. In addition, we also included the interaction of GRS with CxR_t_ (multiplying both variables after they were normalized) to account for potential asymmetric effects of GRS and rewarded and unrewarded trials.

We applied the GLM model to the stay/switch decisions and analyzed the resulting beta weights. To account for outliers, we log-transformed the beta weights and implemented an outlier rejection procedure. We only included sessions whose beta weights were within three standard deviations from the mean. From the above analysis, this led to the exclusion of 6 sessions across 3 monkeys in and a further 35 further human participants. Beta weights were then further submitted to a second-level analysis.

### First level analyses applied to both human and macaque data: Reinforcement learning modelling and bias by irrelevant alternatives

In both species we fitted a reinforcement learning model comprising two model parameters: learning rate and inverse temperature. Specifically, we fit a reinforcement learning model with a Boltzmann action selection rule that uses information about past choices and rewards to estimate expected value for each option on each trial. The reward learning rate and inverse temperature were fitted individually to each session’s data using standard nonlinear minimization procedures. These parameters respectively reflect the weight of influence of the prediction error and the influence of value difference of the probability of choosing an option. To account for outliers in the inverse temperature estimates, we log normalised this parameter. The learning rate and inverse temperature parameters were then submitted to second level analyses.

The reinforcement learning model formed the basis for our investigation of choice biases induced by irrelevant alternatives. This approach investigates whether choices were unduly influenced by the value of an irrelevant alternative. Specifically, we examine whether the interactive impact of the decision-irrelevant option’s value (V_D_) on the choice between the two relevant Options (V_X_ and V_Y_). We applied a two-step multinomial logistic regression analysis, that has been described in full, alongside the complete set of equations, in Noonan et al. (2017). In short, the regression analyses reframe the 3-choice decision as two binary value comparisons between pairs of options. The GLM aims to predict the proportion of choices among the three options from their expected values, with one option assigned in each decision frame as the reference category. For example, Option X and Y are the options being compared; with Option Y as the reference, Option X as the comparator and Option D denoting the irrelevant option. Each option’s values (V_X_, V_Y_, V_D_) is initially derived from a reinforcement learning model described above. The key step, for the purposes of the present study is the isolation of the contextual decision-making factor (V_X_ - V_Y_)V_D_ from the final step of the GLM outlined in equation 1 (equation 7 in Noonan et al (2017)).

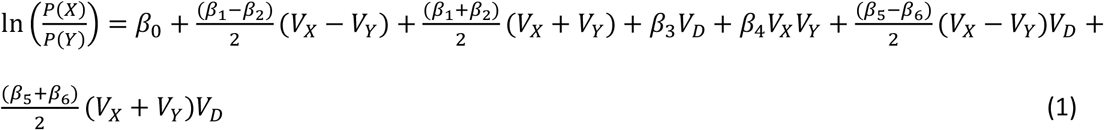

This factor allows us to examine how the expected value of the irrelevant option V_D_ affects the comparison between X and Y (i.e. 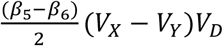), after controlling for the effects of V_X_-V_Y_, V_X_+V_Y_, V_X_×V_Y_, V_D_ and (V_X_+V_Y_)V_D_. In other words, this beta weight reflects the degree to which the effect of value difference between X and Y on choices between these two options was modulated by the irrelevant distractor value (V_D_). For brevity, in the main text, we refer to our variable of interest, the (V_X_ - V_Y_)V_D_, as *bias by irrelevant alternative* (BIA).

In addition to the standard exclusion criteria, the regression model described below failed to fit 6 sessions across from 4 monkeys and 3 sessions from another monkey and a total of 32 human participants these were consequently excluded from the analyses described in this section. The factor isolated from the GLM was subjected to an outlier rejection procedure (6 sessions across 4 monkeys and 15 humans), and the beta weights were absolute log transformed. Beta weights were then submitted to a second-level analysis.

### Second-level analyses in monkeys

Second-level analyses (i.e. averaging over subjects and sessions) were performed in a conceptually similar manner for monkeys and humans, but differed in their implementation because of the nature of the acquired data and the goals of the analyses. For instance, we acquired several sessions worth of data for the same macaques, whereas there was only one session per human participant. Also, in humans, we investigated age-effects whereas in the animals we examined lesion effects.

For both our macaque credit assignment GLM and context decision-making GLM, as well as the derived model parameters (see Supplementary Fig.S1), we submitted resulting (outlier-corrected) beta weights/parameter estimates to separate linear mixed effects models (LME; using Matlab’s fitlme) because the LMEs allowed us to account for monkey identity (‘Mk’) in our analyses. We grouped the data for each second level analysis in three conditions: a baseline condition (all data that were collected in non-lesioned animals), a Lateral lesioned condition (data that were collected from Lateral lesioned animals), and a medial lesioned condition (data that were collected from Medial lesioned animals). All analyses collapsed over experimental paradigms. We coded monkey identity as a random effect with a random intercept and random slopes for all fixed effects used in the LMEs. For significance testing of fixed effects, we used a likelihood ratio test comparing a full model with a model leaving out the particular fixed effect of interest. In addition, we report the fixed effects slope estimates and their standard errors.

The first set of credit assignment LMEs examined deficits in contingent reward assignments after Lateral lesions using a fixed effect of ‘LesionType’ that discriminated between baseline sessions and sessions recorded from Lateral lesioned animals. For this, we examined differences in the effects of the contingent reward history (CxR-history) on staying with the current choice. We considered time points RxC_t_, RxC_t-1_, RxC_t-2_, and RxC_t-3_. For each time point, we constructed an LME as follows (using RxC_t_ as an example) in Wilkinson-Rogers notation:

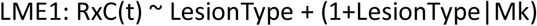

Subsequently, we examined effects of the global reward state (GRS) on choice. To first demonstrate that such effects exist in our data at all, we tested whether the intercept of the LME differed from zero in separate LMEs for each lesion condition. The LMEs comprised only an intercept and the random effect of monkey identity and we compared them with LMEs without intercepts to demonstrate positive GRS effects in both conditions. The LME with intercept was constructed as follows and this procedure was applied to the baseline condition as well as to the lesion condition separately:

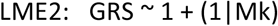

Then, we tested our hypothesis that not only contingent reward assignments but also spread-of-effect related learning mechanisms are impacted by Lateral lesions. To do so, we compared the GRS effects on choice between our baseline data with our Lateral lesion data (fixed effect ‘LesionType’: baseline vs Lateral). We used an LME with the same structure as for the contingent reward assignments described in LME1.

As follow-up comparisons, we tested whether the mechanism by which the GRS impacts reward learning is similarly affected by Medial lesions. To do so, we first demonstrate positive GRS effects in our Medial condition using LME2. Subsequently we ran LME1 twice, once to determine differences between the baseline condition and the Medial condition (LesionType: baseline vs Medial) and once to determine differences in effect sizes between the two lesion conditions themselves (LesionType: Medial vs Lateral).

We followed a similar LME analysis structure for the analysis of the critical factor derived from the contextual decision-making GLM: the BIA effect. For the latter variable we examined the degree of influence each factor had on X/Y choices. We tested for deficits in context-based decisions after Medial lesions using LME1 with a fixed effect of ‘LesionType’ that discriminated between baseline sessions and sessions taken from Medial lesioned animals. We performed the equivalent LMEs for the Lateral lesioned animals, using LME1 to examine fixed effects of ‘LesionType’ (LesionType: baseline vs Lateral).

### Second-level analyses in humans and human specific analyses

Second-level analyses in our human data focused on age-related differences in first-level effect sizes. Because there is some uncertainty about the precise age when development cognitive changes in credit assignment are expected to occur and because it is unclear if these changes are expected to happen in a continuous fashion or step-wise, we analysed age effects in our human experiment in two complementary analysis ways. All our key results survive both ways of analyzing age effects. First, we performed an age split and compared an adolescent sub-group (age < 18; n = 159) with a group of young adults (age ≥ 18; n = 228) via independent samples t-tests (Fig.3C). Should developmental changes in computational subprocesses of reward learning occur during adolescence, then we should expect significant differences between the two groups. However, we also analyzed our data in a continuous way as cognitive processes may slowly mature over time irrespective of precise age boundaries. For this, we used Pearson’s linear correlation analyses with age. After demonstrating that such a correlation exists for our variables of interest, we then fitted linear, quadratic and cubic link functions. The purpose of these follow-up analyses was to develop a more detailed picture of the maturation of the variable of interest over our entire age range of 11 to 35 years (Xia *et al*., 2020). Among these functions, we identified the one with the best fit as indicated by the lowest AIC value. In addition, we ran a likelihood-ratio, using the corresponding models least-squares error, to confirm that the best fitting model explained significantly more variance than a null model. We constructed the null model by assuming no relationship with age and instead fitted the mean of the observed dependent measures. As summary statistics we report Pearson’s R for correlations with age, the *p* value for likelihood-ratio test (LRT) and the adjusted R^2^ of the best fitting model.

First, we performed two unique additional analyses in humans to broadly detail our human data. As initial analyses of task performance, for each participant calculated total rewards earned during the task. Next, we calculated the proportion of best. This measure was derived from estimated expected value of each stimulus option using the reinforcement learning model as described above. Both metrics were subjected to the two-step pipeline described for second level analyses in humans.

Next, we used the two-step analysis pipeline for the human credit assignment analysis. We analysed [1] the CxR_t_ within the CxR-history component, [2] the C_t-1_ factor within the C-history component (see Supplementary Fig.S2) and [3] the single GRS factor. To complement the GRS analyses described above for both species, and in line with the analysis approach described previously (Wittmann *et al*., 2020), we conducted a follow-up analysis in our human data to investigate the influence of the GRS on switch/stay decisions. We estimated the residual probabilities of a choice to switch or stay by regressing out of the data all effects of the previous GLM, except CxR_t_ and GRS. The resulting choice residual were then binned by [1] the receipt of a reward on trial t, and [2] GRS (low or high; calculated as a median split of GRS). The estimated residual probabilities derived from the subsidiary GLM investigating the influence of the GRS on switch/stay decisions were split into adolescents and adult age categories and subjected to a 2 (receipt of reward on trial t [reward; no reward]) x 2 (GRS [low; high]) x 2 (age[adolescent; adult]) repeated measures ANOVA. According to the outlier rejection procedure described above, now only a single participant was excluded from this follow-up analysis

In humans, we also examined a linear relationship between the GRS effects and contingent learning in humans. For this, we calculated a correlation between CxR_t_ and GRS, which reflect key markers of contingent credit assignment and GRS effects, respectively. As this analysis was aimed to show age-general relationship, we calculated a partial correlation between the two variables controlling for age and the GLM constant. This was to ensure this finding would not be confounded by age differences and the baseline tendency of participants to stay or switch.

Finally, in humans, we examined age-related differences in model fit parameters, learning rate and inverse temperature, and the contextual decision-making factor, BIA. These model parameters were all compared across age using the two-step process outlined above.

## Acknowledgements

The authors would like to acknowledge the early contributions to the study made by Juliette Westbrook, Morwenna Rickards and Linnet Chan as part of their 3^rd^ year project. We would also like to thank Matthew Rushworth for his constructive feedback on early versions of the manuscript. We are also grateful to all our participants, young and older for contributing to the study.

