## Supplementary Information for "Dissociable mechanisms of reward learning co-mature during human adolescence as predicted by macaque lesion models"

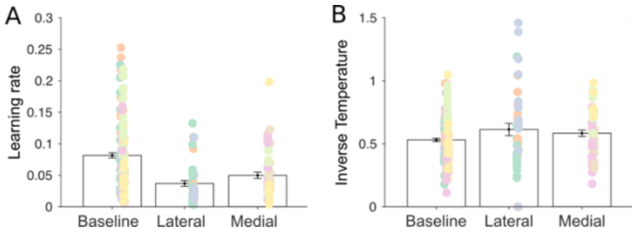


**Figure S1. Learning rate, but not choice accuracy, are affected by orbitofrontal lesions.** We examined the fitted parameters, learning rate and inverse temperature, of the reinforcement learning model fit to monkeys choices with LMEs testing between-group differences after either lateral (LesionType: baseline vs Lateral) or medial (LesionType: baseline vs Medial) orbitofrontal lesions. Reward learning rate, coded in the anterior cingulate cortex (Behrens *et al.*, 2007), is a parameter that reflects the impact that a single outcome has on the value of the chosen option and is considered conceptually related to contingency learning. Mirroring the contingency learning deficits described after Lateral lesions, these animals also showed lower learning rates than animals in the baseline conditions (base v Lateral: Estimate=-0.005, SE=0.001, X2(1)=1, p = 0.008). However, the learning rate may actually be a more general learning mechanism as we observed that Medial lesioned animals also showed reduced learning rates (base vs Medial: Estimate=-0.003, SE=0.001, X2(1)=1, p=0.029). These findings are notably independent of changes in the animals' general levels of decision-making noise, with the RL models' inverse temperature not affected by either lesion (base v Medial: Estimate=-0.055, SE=0.039, X2(1)=1, p=0.183, base v Lateral: Estimate=0.055, SE=0.048, X2(1)=1, p=0.305).

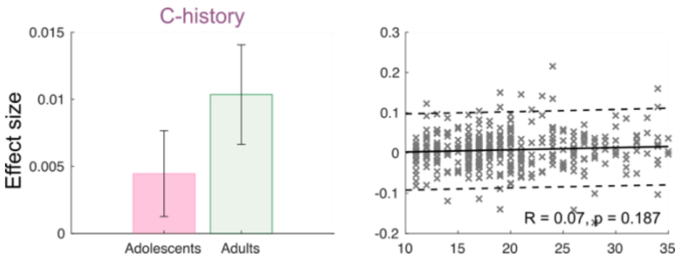


**Figure S2. Influence of choice history on switch/stay decisions does not change across adolescence.** As a control analysis, we also considered developmental changes in a reward-unrelated learning mechanism that is not linked to orbitofrontal cortex and that was also included in our credit assignment GLM. We examined C-history, the tendency to repeat choices irrespective of reward (Lau & Glimcher, 2005; Akaishi *et al.*, 2014; Wittmann *et al.*, 2020). In general, participants were more likely to repeat the most recent choice, irrespective of reward (not shown, one-sample t-test; $C_{t-1}$: $t_{352}$=3.09, p=0.002). However, C-history did not differ between adolescents and adults ($t_{351}$=1.16, p=0.245) and there was no correlation between age and $C_{t-1}$ (R=0.07, p=0.191). Note that these effects were not only not significant, they were also in the direction contrary to the idea that choice repetition decreases over age. Note that such a decrease would have been a plausible alternative by which task performance could have improved, as choice repetition per se may lead participants to discount appropriate choice-reward contingencies.
